# The *C. elegans* LON-1 protein requires its CAP domain for function in regulating body size and BMP signaling

**DOI:** 10.1101/2024.10.01.616164

**Authors:** Maria Victoria Serrano, Stephanie Cottier, Lianzijun Wang, Sergio Moreira-Antepara, Anthony Nzessi, Zhiyu Liu, Byron Williams, Myeongwoo Lee, Roger Schneiter, Jun Liu

## Abstract

The CAP (Cysteine-rich secretory proteins, Antigen-5, Pathogenesis-Related) proteins are widely expressed and have been implicated to play diverse roles ranging from mammalian reproduction to plant immune response. Increasing evidence supports a role of CAP proteins in lipid binding. The *C. elegans* CAP protein LON-1 is known to regulate body size and Bone Morphogenetic Protein (BMP) signaling. LON-1 is a secreted protein with a conserved CAP domain and a C-terminal unstructured domain with no homology to other proteins. In this study, we report that the C-Terminal Domain (CTD) of LON-1 is dispensable for its function. Instead, key conserved residues located in the CAP domain are critical for LON-1 function in vivo. We further showed that LON-1 is capable of binding sterol, but not fatty acid, in vitro, and that certain key residues implicated in LON-1 function in vivo are also important for LON-1 sterol binding in vitro. These findings suggest a role of LON-1 in regulating body size and BMP signaling via sterol binding.

**ARTICLE SUMMARY:** The *C. elegans* LON-1 protein is known to regulate body size and Bone Morphogenetic Protein (BMP) signaling. However, its molecular mode of action remains elusive. This study shows that LON-1 can bind sterol, but not fatty acid, in vitro. Furthermore, key conserved residues in the CAP domain of LON-1 are required for LON-1 function in vivo. These findings suggest a role of LON-1 in regulating body size and BMP signaling via sterol binding.

## INTRODUCTION

Bone Morphogenetic Proteins, or BMPs, are a class of signaling molecules in the TGFβ superfamily. BMP signaling molecules are conserved across metazoans with well-established roles in development and homeostasis. Disruptions of this pathway in humans are associated with a broad range of disorders, including cancers, musculoskeletal abnormalities, and other adverse health outcomes (WANG *et al*. 2014; KATAGIRI AND WATABE 2016; AKIYAMA *et al*. 2024). Thus, BMP signaling needs to be tightly regulated to ensure spatiotemporal specificity with the optimal outcome.

The nematode *C. elegans* provides a unique system for dissecting mechanisms of BMP signaling output and its regulation. All the core BMP signal transduction pathway members are conserved in *C. elegans,* yet BMP signaling is not essential for *C. elegans* viability or fertility. Instead, BMP signaling is known to regulate body size and mesoderm development, as well as a number of other processes (SAVAGE-DUNN AND PADGETT 2017). Increased BMP signaling leads to a long body size, while decreased BMP signaling causes a small body size phenotype. In addition, mutations in the zinc finger transcription factor SMA-9, a Schnurri homolog, cause a dorsal-to-ventral fate transformation in the *C. elegans* postembryonic mesoderm (LIANG *et al*. 2003; FOEHR *et al*. 2006). Mutations in core members of the BMP pathway can specifically suppress this mesodermal defect of *sma-9(0)* mutants (FOEHR *et al*. 2006; LIU *et al*. 2015). Importantly, mutations in factors involved in modulating the BMP pathway can also suppress the *sma-9(0)* mesoderm defect (TIAN *et al*. 2010; TIAN *et al*. 2013; LIU *et al*. 2015; WANG *et al*. 2017; DEGROOT *et al*. 2019).

Using a screen for mutations that can suppress the *sma-9(0)* mesoderm phenotype, we have previously identified LON-1 as a modulator of the BMP pathway (LIU *et al*. 2015). LON-1 is known as a negative regulator of body size, as *lon-1* mutant alleles produce worms with a long body length, and that *lon-1* transcription is negatively regulated by the BMP signaling pathway (MADUZIA *et al*. 2002; MORITA *et al*. 2002). However, how LON-1 acts to regulate body size and modulate BMP signaling is not well understood. In this study, we carried out structure-function analysis of LON-1, with the goal of understanding its mechanism of action.

LON-1 is a predicted secreted protein belonging to the CAP protein superfamily, with an additional C-terminal tail that is predicted to be unstructured (Figure 1). CAP (Cysteine-rich secretory proteins, Antigen-5, Pathogenesis-Related 1) superfamily (pfam PF00188) members are present in every kingdom of life, and they share a conserved CAP domain of approximately 150 amino acids that forms a characteristic alpha-beta-alpha structure (GIBBS *et al*. 2008; SCHNEITER AND DI PIETRO 2013; ABRAHAM AND CHANDLER 2017; GAIKWAD *et al*. 2020) (Figure 1C,D). CAP proteins are widely expressed and have been found to have diverse roles ranging from mammalian reproduction to plant immune response and venom toxicology (GIBBS *et al*. 2008; GAIKWAD *et al*. 2020). One of the best studied CAP proteins is yeast Pry1. Pry1 is known to transport sterols and fatty acids out of the cell (CHOUDHARY AND SCHNEITER 2012; DARWICHE *et al*. 2016; DARWICHE *et al*. 2017). Moreover, mutational studies showed that Pry1 binds to fatty acid and sterol via independent sites within the CAP domain (DARWICHE *et al*. 2017). This small molecule binding activity of the CAP domain seems to be conserved despite the diversity of the CAP superfamily, as human CRISP2, for example, can bind cholesterol in vitro and can rescue the cholesterol accumulation phenotype seen in Pry1 deficient yeast (CHOUDHARY AND SCHNEITER 2012). More recently, two CAP domain proteins in *C. elegans*, SCL-12 and SCL-13, were found to bind to cholesterol and play a role in the mobilization of cholesterol during the transition of dauer worms from quiescence to growth (SCHMEISSER *et al*. 2024).

**Figure 1.**
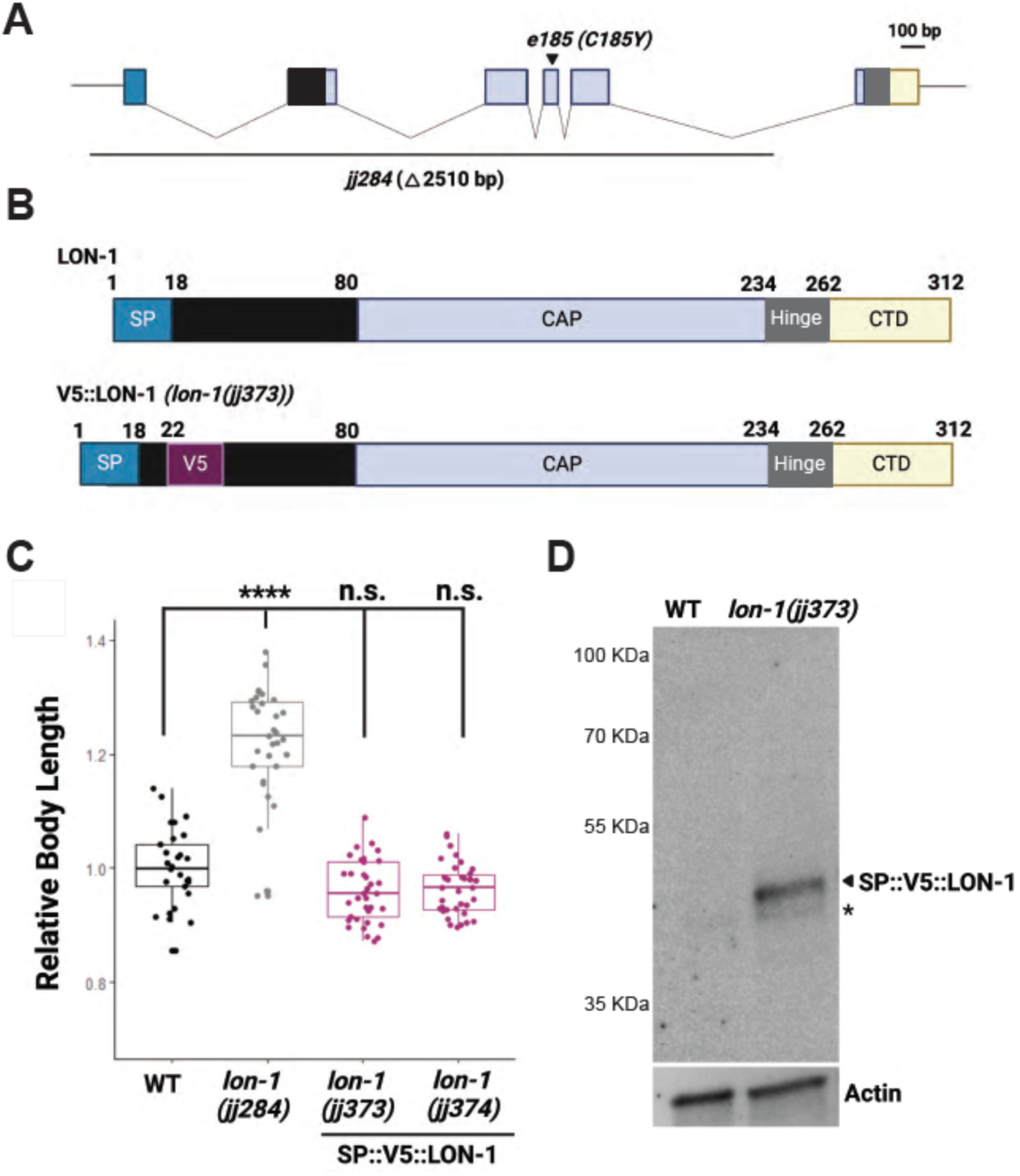
Summary of null and endogenously tagged *lon-1* alleles. (A) Schematic of the *lon-1* gene, including the canonical *e185* allele, and the null allele generated in this study, *jj284*. (B) Schematics of WT LON-1 protein and tagged SP::V5::LON-1 protein present in *lon-1(jj373)* animals. LON-1 contains a predicted SP (signal peptide), a CAP (Cysteine-Rich secretory protein, Antigen-5, Pathogenesis-Related 1) domain, a Hinge region, and an unstructured CTD (C-terminal Domain) with no known homology. (C) Relative body lengths of *lon-1* alleles at larval L4.3 stage, normalized to WT. Each dot represents one worm. **** *p*< 0.0001; n.s. not significant. Tested using an ANOVA with a Tukey HSD. (D) Western blot showing that SP::V5::LON-1 protein is detectable using anti-V5 (top). Actin was used as a loading control (bottom). Each lane contains lysates from 100 young adult worms.

Many CAP family members also contain an additional C-terminal domain of distinct types after the CAP domain (ABRAHAM AND CHANDLER 2017; GAIKWAD *et al*. 2020). These additional domains are thought to confer specificity to CAP protein functions, such as the ion channel regulatory domain (ICR) within the CRISP (Cysteine-rich secretory protein) subfamily of CAP proteins that confers ion channel blocking activities (GAIKWAD *et al*. 2020). In this study, however, we report that the C-terminal domain (CTD) of LON-1 is dispensable for its function. Instead, key conserved residues located in the CAP domain are critical for LON-1 function in regulating body size and BMP signaling. Moreover, LON-1 is capable of binding sterol in vitro. These findings suggest that the CAP protein LON-1 may also function by binding sterols in regulating body size and BMP signaling.

## MATERIALS AND METHODS

### C. elegans strains

All strains (Supplemental Table 1) used in this study were maintained using standard culture methods as described in (BRENNER 1974). Strains were maintained at 20°C unless otherwise specified.

### Generation of lon-1 alleles via CRISPR

Both the *lon-1* deletion allele *jj284* and the endogenously tagged SP::V5::LON-1 alleles, *lon-1(jj373)* and *lon-1(jj374),* were generated via CRISPR by injecting plasmids expressing Cas9 (pDD162, (DICKINSON *et al*. 2013) and the appropriate sgRNA(s) in the pRB1017 backbone (ARRIBERE *et al*. 2014), the single stranded homologous repair template (Supplemental Table 2). PRF4(*rol-6(d)*) (MELLO *et al*. 1991b) was used as a co-injection marker. The guide RNA plasmids for generating SP::V5::LON-1 are pZL96 and pZL97, while the guide RNA plasmids for generating *lon-1(jj284)* are pZL96, pZL97 and pZL99 (Supplemental Table 3). One allele, *lon-1(jj373)*, underwent repair as expected and contains a 9 aa linker (GASGSSGAS) between the inserted V5 tag and the remainder of the LON-1 sequence. The second allele, *lon-1(jj374)*, contains a mutation in the linker that instead renders the sequence GASGSSEAS. Both alleles behaved identically.

To create specific point mutations in *lon-1*, CRISPR target sites were identified within the gene using SnapGene software (www.snapgene.com). The *lon-1* mutant alleles, such as *kq153*, *kq171*, *kq175*, *kq185,* and *kq207*, were generated by standard microinjection and PCR genotyping. Briefly, the mixture of custom repair template oligos (Supplemental Table 1), custom guide RNAs, tracrRNA (cat. #1072532, IDT Inc.), and Alt-R Cas9 nuclease (cat. #1081058, IDT Inc.) was annealed at room temperature (PAIX *et al*. 2015) and micro-injected into the syncytial gonad arms of N2 wild-type or LW6100[*lon-1(jj373[sp::v5::lon-1])*] worms (P0) (MELLO *et al*. 1991a; MELLO *et al*. 1991b). The injection mix also contained *dpy-10* guide RNA (ZQDP10A, Supplemental Table 2) as a co-CRISPR marker (ARRIBERE *et al*. 2014). The F1 animals were screened for the intended mutation by identifying animals exhibiting the Rol phenotype and PCR genotyping using the mutation-specific primers (Supplemental Table 1). The PCR genotyping was repeated on the non-roller F2 offspring from the F1 animals carrying the mutation using wild-type (WF) and mutation-specific (MF) primers (Supplemental Table 1) to isolate homozygous mutants. The F2 animals showing only the mutation-specific PCR product were considered homozygous. Similar methods were used to generate the *lon-1* CTD deletion alleles *lon-1(jj373 jj536)* and *lon-1(jj373 jj537)*.

All CRISPR alleles were confirmed by Sanger sequencing, and outcrossed with wild-type worms for at least two rounds before further analyses.

### Body length measurement

Worms at early larval L4 stage were selected from mixed-stage plates and immobilized in a 0.35 mM levamisole in M9 solution. Substages were classified based on vulval development as described in Mok et al (MOK *et al*. 2015). Worms were mounted onto glass slides with 2% agarose pads, and imaged using a Leica DMRA2 compound microscope equipped with a Hamamatsu Orca-ER camera and the iVision software (Biovision Technology). Images were taken using the 10x objective and body length of worms was determined using the segmented line tool in Fiji, following the midline of the worm from head to tail. Statistical analysis was conducted using Tukey’s Honest Significant Difference (HSD) tool in Rstudio.

### sma-9(0) suppression (Susm) assay

To determine the penetrance of the Susm (suppression of *sma-9(0)* M-lineage defect) phenotype, worms were grown at 20°C and scored as adults for the number of coelomocytes (CCs) using a secreted *CC::GFP* marker, *arIs37* (FARES AND GREENWALD 2001). The number of worms with 6 CCs was divided by the total number of worms scored to generate the Susm penetrance. For each genotype, 2 independent isolates were generated (Supplemental Table 1), and 4 to 6 plates of worms from each isolate were scored for the Susm penetrance. The Susm data from 2 isolates were combined and the means of the Susm penetrance were presented. Statistical test was done using an unpaired two-tailed student *t* test.

### Drosophila S2 cell culture and transfection

*Drosophila* S2 cells were grown in M3+BPYE+10% Heat-inactivated FBS and transfected using a calcium phosphate method (Invitrogen protocol, Version F 050202 28-0172). Cells and their corresponding media were collected 7 days post-transfection. Cells were separated from media by centrifugation (4K, 1 minute). Cell pellet was washed twice in PBS and then lysed in 300 ul lysis buffer (50 mM Tris pH 7.6, 150 mM NaCl, 1 mM EDTA, 1% Triton-X). Protease inhibitor (Pierce, A32965) was added to all samples to avoid protein degradation. To prepare samples for western blotting, 20 ul of total cell lysate was mixed with 5 ul of 5xSDS sample buffer (0.2 M Tris-HCl (pH 6.8), 20% glycerol, 10% SDS, 0.25% bromophenol blue, 10% β-mercaptoethanol) and heated at 95°C for 5 minutes. 10 ul of this mixture was used as a sample for western blotting. For cell media, 10 ul was taken from the 4 mL of media generated by the above protocol and mixed with 2.5 ul of 5xSDS sample buffer (0.2 M Tris-HCl (pH 6.8), 20% glycerol, 10% SDS, 0.25% bromophenol blue, 10% β-mercaptoethanol) for use as a sample for western blotting.

### Gel electrophoresis and western blotting

One hundred young adult or gravid adult worms were collected in 20 ul H_2_O and mixed with 5 ul of 5xSDS sample buffer (0.2 M Tris-HCl (pH 6.8), 20% glycerol, 10% SDS, 0.25% bromophenol blue, 10% β-mercaptoethanol) before flash freezing in liquid nitrogen and then immediately heating at 95°C for 10 minutes. Samples were loaded onto 10% Mini-PROTEAN TGX Precast Gels (Bio-Rad Laboratories) at 150 V and run using the Biorad Mini Protean Tetra system and transferred for 7 minutes to Immobilon-P PVDF membrane (MilliporeSigma) using an Invitrogen semi-dry Power Blotter Station. Membranes were blocked for 1 hour at room temperature in 10% milk (Carnation) in 1x PBST (137 mM NaCl, 2.7 mM KCl, 10 mM Na2HPO4, 1.8 mM KH2PO4, 0.1% Tween-20). Membranes were incubated with primary antibodies in 5% milk in 1xPBST at 4°C overnight, and with secondary antibodies in 5% milk in 1xPBST for 2 hrs at room temperature. Primary antibodies used include: rabbit anti-V5 Ab (Cell Signaling Technology Product Number D3H8Q), mouse anti-Actin Ab (1:2000, Developmental Studies Hybridoma Bank Product ID JLA20). Secondary antibodies used include: HRP-conjugated Goat anti Rabbit IgG (Jackson ImmunoResearch Laboratories Inc, 1:10000) and IRDye 800CW conjugated goat anti-mouse IgM (LI-COR, 1:5000). Membranes were incubated with BioRad Clarity Western ECL substrate for 5 minutes before imaging using a BioRad ChemiDoc MP imaging system.

### Plasmid preparation for protein isolation

*lon-1* ORF with codons optimized for expression in *Escherichia coli* was synthesized by GenScript (Rijswijk). Its coding sequence devoid of the signal sequence (starting at Glycine 24) was amplified by PCR using KAPA HiFi DNA Polymerase (Roche) and cloned into BamHI/XhoI digested pET28a-6xHis-SUMO vector using Gibson Assembly® Master Mix (New England Biolabs). The *lon-1* mutations E153A, V175A, C185Y and C207S were introduced by PCR-ligation-PCR approach (ALI AND STEINKASSERER 1995), and then the PCR products were cloned with the same strategy as for wild-type *lon-1*. All constructs were verified by Sanger sequencing (Microsynth AG).

### Protein expression and purification

LON-1 proteins (wild-type and mutants) were expressed in BL21 (DE3) *E. coli* cells (Novagen). Transformed bacteria were grown at 37°C in LB medium supplemented with Kanamycin until OD_600_ reached 0.5. The cells were then shifted to 16°C and protein expression was induced over night by addition of lactose (50 mM). Cells were harvested, resuspended in lysis buffer (50 mM Tris-HCl pH 7.5, 300 mM NaCl, 20 mM imidazole, 1 mM PMSF, 1x EDTA-free protease inhibitor cocktail (Roche) and broken open using a Microfluidizer LM10 (Microfluidics). Soluble proteins were incubated with nickel-nitriloacetic acid beads (Qiagen) for 2 h at 4°C. Bound proteins were then washed, and eluted with lysis buffer containing 300 mM imidazole and 10% glycerol. Finally, the proteins were loaded on Zeba spin desalting columns (ThermoFisher Scientific) to remove imidazole by buffer exchange (50 mM Tris-HCl pH 7.5, 300 mM NaCl, 1 mM PMSF, 1x EDTA-free protease inhibitor cocktail, 10% glycerol). Protein concentration was determined using the Qubit^TM^ Protein and Protein Broad Range (BR) Assay Kit (Invitrogen).

### In vitro lipid binding assays

Binding of wild-type and LON-1 mutant proteins to cholesterol sulfate (stock solution: 10 mM DMSO, Sigma) and palmitic acid (stock solution: 78 mM ethanol, Sigma) was assessed by microscale thermophoresis (MST) using a Monolith NT.115 system (Nanotemper Technologies) in binding buffers consisting of 200 mM Bis-Tris pH 6.5, 50 mM NaCl, 0.1% Triton X-100, and 50 mM Tris-HCl pH 7.5, 30 mM NaCl, 0.05% Triton X-100, respectively. Briefly, purified proteins were labeled with His-Tag labeling kit RED-tris-NTA following the manufacturer’s instructions (Nanotemper technologies), and 50 ng of labelled proteins were mixed with serial dilution of the lipids of interest and loaded into MST standard capillaries. Using the KD Model of the MO.Affinity Analysis software (Nanotemper Technologies), dissociation constants were obtained by plotting fraction-bound against the logarithm of ligand concentration. The data were exported and plotted in Prism 9. Experiments were performed in triplicates.

## RESULTS AND DISCUSSION

### Generation of a molecular null allele of *lon-1* and a strain that expresses a fully functional, endogenously tagged LON-1

As *lon-1* alleles used in previous studies were all point mutations that either cause single amino acid changes or pre-mature stop codons (MADUZIA *et al*. 2002; MORITA *et al*. 2002), we first used CRISPR to generate a molecular null allele of *lon-1*. This new allele, *lon-1(jj284),* contains a 2510 bp deletion, beginning 220 bp upstream of the ATG and ending 281 bp into the last intron (Figure 1A). *lon-1(j284)* worms exhibit a long body length and a *sma-9(0)* suppression (Susm) penetrance of 45% (Figure 1C, Table 1), consistent with previously reported roles of LON-1 in the regulation of body size and BMP signaling (MADUZIA *et al*. 2002; MORITA *et al*. 2002; LIU *et al*. 2015).

**Table 1.** Summary of the Susm phenotype of various *lon-1* alleles.

| Genotype | Susm penetrance | N |
| --- | --- | --- |
| *sma-9(cc604)* | 0% | 709 |
| *lon-1(jj284); sma-9(cc604)* | 45%*** | 1473 |
| *lon-1(e185); sma-9(cc604)* | 21%*** | 414 |
| *lon-1(jj373[SP::V5::LON-1]); sma-9(cc604)* | 0.6%<sup>n.s.</sup> | 502 |
| *lon-1(jj374[SP::V5::LON-1]); sma-9(cc604)* | 0.6%<sup>n.s.</sup> | 318 |
| *lon-1(jj373 jj536[SP::V5::LON-1(ΔCTD)]; sma-9(cc604)* | 1.2%<sup>n.s.</sup> | 584 |
| *lon-1(jj373 jj537[SP::V5::LON-1(ΔCTD)]; sma-9(cc604)* | 1.4%<sup>n.s.</sup> | 275 |

Susm penetrance was measured as the percentage of 6 CC animals as scored by the *arls37(secreted CC::GFP)* reporter. Statistical analysis was conducted by comparing double mutant lines with the *sma-9(cc604)* single mutants. \*\*\* $p < 0.001$ ; n.s. not significant (unpaired two-tailed Student's t test).

We then used CRISPR to insert the V5 tag to the LON-1 N-terminus following the signal peptide (Figure 1B). We isolated two independent alleles, *lon-1(jj373[SP::V5::LON-1])* and *lon-1(jj374[SP::V5::LON-1])* (see Materials and Methods). Both alleles exhibited the WT body length and an absence of the Susm phenotype (Figure 1C, Table 1), suggesting that the tagged LON-1 protein is fully functional. The SP::V5::LON-1 protein is detectable via western blot at the predicted molecular weight (Figure 1D), providing us with a tool to monitor LON-1 protein in vivo.

### LON-1 is a secreted protein in the CAP superfamily

The N-terminus of the LON-1 protein contains a stretch of hydrophobic residues, which could function as a signal peptide (Figure 1A; as suggested by (MADUZIA *et al*. 2002) and supported by programs that predict signal peptides) or as a transmembrane domain (as suggested by (MORITA *et al*. 2002)). To resolve this issue, we expressed in *Drosophila* S2 cells full length LON-1 tagged with the V5 epitope at its C-terminus (Figure 2B). We could detect LON-1 protein in the culture medium of transfected cells (Figure 2C), suggesting that LON-1 can be secreted and that the N-terminal hydrophobic stretch in LON-1 likely acts as a signal peptide for secretion. Despite being secreted, LON-1 may have some affinity to the membrane (as observed by (MORITA *et al*. 2002). This is not unprecedented, as the secreted *Candida albicans* CAP protein Rbe1p associates with the cell wall in a disulfide-dependent manner (BANTEL *et al*. 2018).

**Figure 2.**
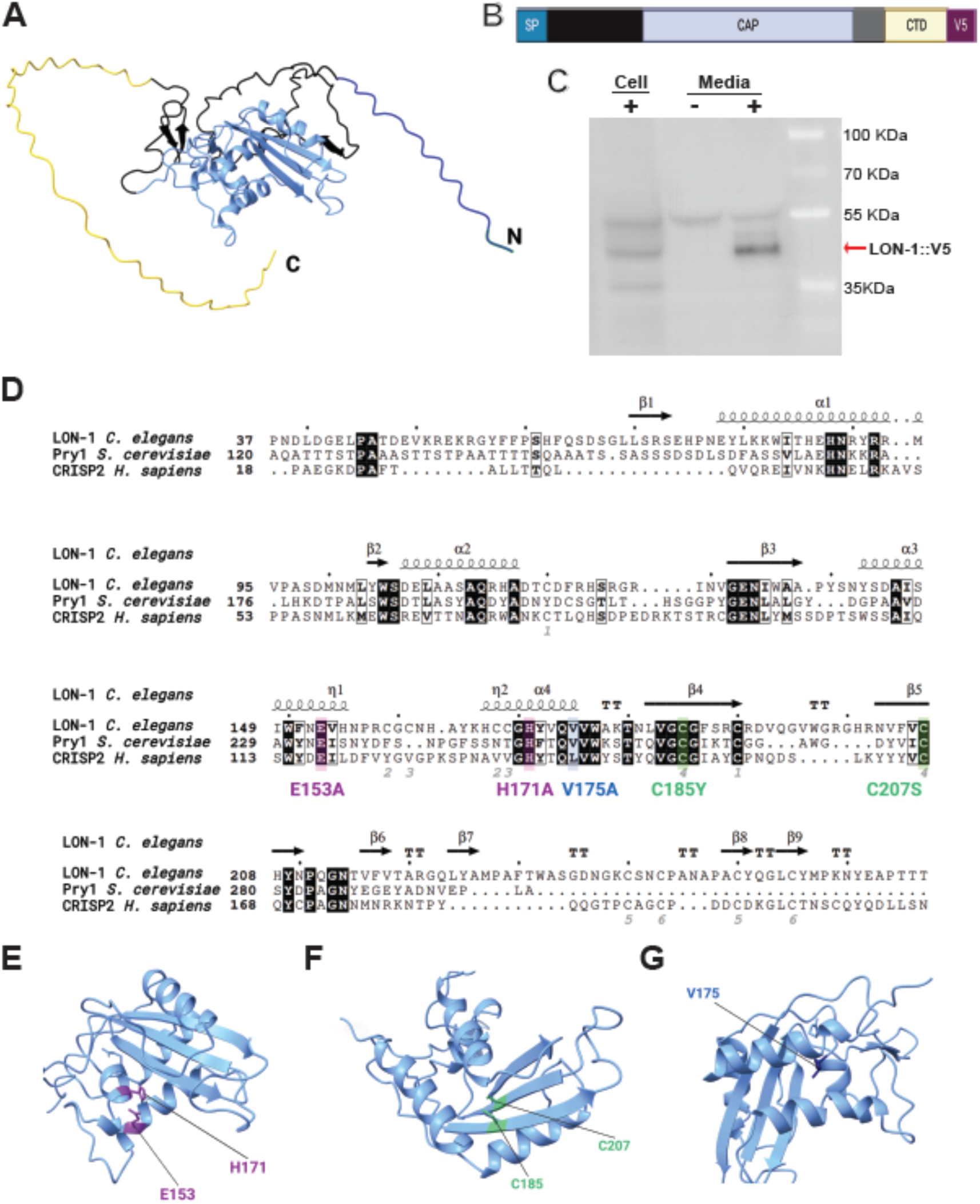
LON-1 is a secreted protein in the CAP superfamily. (A) Predicted structure of LON-1 protein by Alphafold (Model AF-Q09566-F1), visualized using UCSF ChimeraX (MENG *et al*. 2023). The signal peptide is colored in dark blue, the CAP domain in light blue, the Hinge region in gray, and the CTD in yellow. (B) Schematic of the LON-1::V5 protein expressed in *Drosophila* S2 cells. (C) Western blot showing that LON-1::V5 protein is secreted into the media when expressed in *Drosophila* S2 cells. Cells were separated from their corresponding media 7 days after transfection. “+” samples from cells transfected with pAc5::LON-1::V5. “-“: samples from un-transfected cells. (D) Alignment of the CAP domains of LON-1 and two other CAP family members. From top to bottom: *C. elegans* LON-1 (Gene ID: 175753), *Saccharomyces cerevisiae* Pry-1 (Gene ID: 853366), and *Homo sapiens* CRISP2 (Gene ID: 7180). Protein sequence alignment was generated using Clustal Omega, accessed online (https://www.ebi.ac.uk/jdispatcher/msa/clustalo). This figure was generated with ENDscript 2 (ROBERT AND GOUET 2014), using the predicted structure of LON-1 protein by Alphafold (Model AF-Q09566-F1). Color shaded boxes mark conserved amino acids that are mutated in various *C. elegans* point mutants. Mutations in purple (E153A, H171A) target a putative catalytic site, those in green (C185Y, C207S) target a cysteine bridge, and a mutation in blue (V175A) targets a putative fatty-acid binding region. (E-G) Zoomed regions of predicted LON-1 structure showing the locations of amino acids of interest.

Following the signal peptide and beginning about 80 aa into the LON-1 protein sequence is a predicted CAP domain (Figure 1B, 2A,D). The CAP domain is predicted to fold into an α-β-α secondary structure. Sequence alignment and structural prediction using AlphaFold (Figure 2D) indicated that LON-1 shares with other CAP domain proteins a similar structure and conserved key amino acids. For example, two conserved residues in LON-1, H171 and E153, represent two of the four residues referred to as the CAP tetrad, which has been shown in several CAP protein structures to form a divalent cation binding pocket (WANG *et al*. 2010; ASOJO 2011; MASON *et al*. 2014) (Figure 2E). Mg^2+^ binding of the yeast protein Pry1 via this pocket is necessary for the sterol-binding activity of Pry1 in vitro (DARWICHE *et al*. 2016). These observations suggest that if H171 and E153 may be important for LON-1 function in *C. elegans*. Similarly, two cysteine residues C185 and C207 are predicted to form a disulfide bridge in the central beta sheet of the CAP domain (Figure 2F), like other CAP domain proteins (ASOJO 2011; MASON *et al*. 2014). Mutating one of the equivalent residues in yeast Pry1 at C279 disrupted the sterol-binding ability of Pry1 (CHOUDHARY AND SCHNEITER 2012). A similar mutation in the *Solanum lycopersicum* (tomato) PR-1 (Pathogenesis-Related 1) protein P14c also affected the protein’s ability to bind sterol (GAMIR *et al*. 2017). Finally, it has been found that a valine at position 227 in Pry1, together with the second alpha helix, forms a channel responsible for binding fatty acids but not for binding sterol (DARWICHE *et al*. 2017). This valine is conserved in LON-1 as V175 (Figure 2G). Some of these key residues in the CAP domain were found to be important for LON-1 function in *C. elegans* (see below).

### The C-terminal domain (CTD) of LON-1 is not essential for LON-1 function in vivo

In addition to the signal peptide and the CAP domain, LON-1 has a 50 amino acid-long, C-terminal, unstructured and disordered domain with no homology to any other proteins (Figure 3A, Supplemental Figure 1). Not all CAP domain proteins have a C-terminal domain (CTD) (ABRAHAM AND CHANDLER 2017; GAIKWAD *et al*. 2020). But the CAP domain proteins of the CRISP subfamily found in organisms such as insects, mollusks and vertebrates have a C-terminal ion channel regulatory (ICR) domain connected to the CAP domain via a hinge region (ABRAHAM AND CHANDLER 2017; GAIKWAD *et al*. 2020). The LON-1 CTD sequence does not contain the conserved cysteine residues fundamental to the ICR domain of CRISP proteins, but LON-1 does have a hinge region (Supplemental Figure 1). Intriguingly, among members of the CAP domain protein family in *C. elegans*, LON-1 is unusual in having a long CTD, whereas the majority of the other CAP domain proteins end abruptly after the hinge region (Supplemental Figure 1). We therefore tested whether LON-1 requires its CTD for its function in regulating body size and BMP signaling.

**Figure 3.**
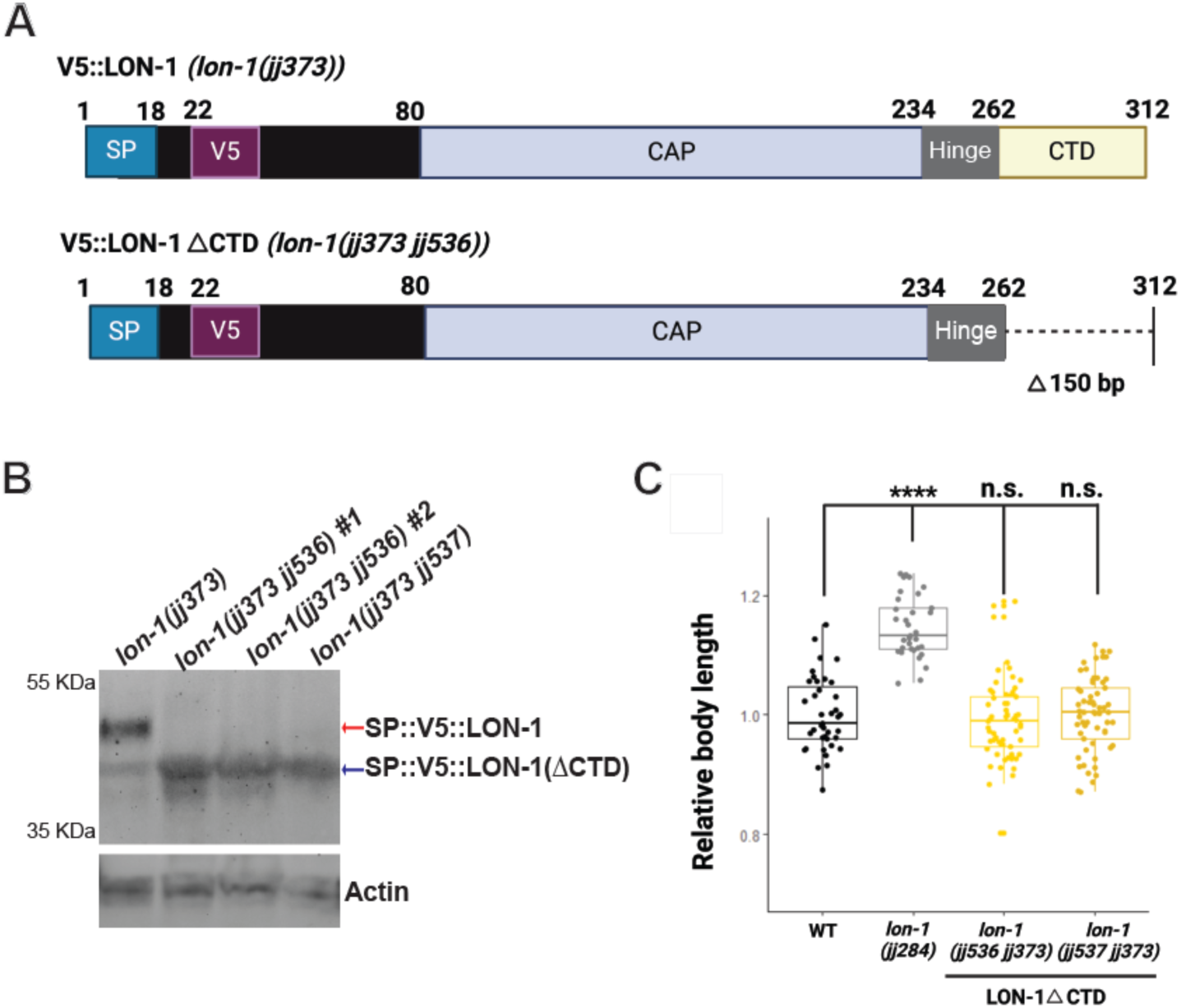
LON-1 CTD is not crucial for LON-1 function in vivo. (A) Schematics of SP::V5::LON-1 present in *lon-1(jj373)* and SP::V5::LON-1**Δ**CTD present in *lon-1(jj373 jj536)* animals. (B) Western blot showing that SP::V5::LON-1(ΔCTD) protein is detectable using anti-V5 (top). Actin was used as a loading control (bottom). Each lane contains lysates from 150 gravid adult worms. The SP::V5::LON-1(ΔCTD) protein is of similar molecular weight to a non-specific protein band (marked with an asterisk in Figure 1D) found in *lon-1(jj373[SP::V5::LON-1])* worms. (C) Relative body lengths of *lon-1* alleles at larval L4.1 stage, normalized to WT. Each dot represents one worm. **** *p*< 0.0001; n.s. not significant. Tested using an ANOVA with a Tukey HSD.

To assess the importance of the CTD in LON-1 function, we used CRISPR to generate a deletion of the last 50 amino acids of LON-1 in the *lon-1(jj373[SP::V5::LON-1])* background. We obtained two deletion alleles, *lon-1(jj373 jj536[SP::V5::LON-1(ΔCTD)])* and *lon-1(jj373 jj536[SP::V5::LON-1(ΔCTD)]). lon-1(jj373 jj536)* is a deletion of 50 amino acids as expected, while *lon-1(jj373 jj537)* contains a deletion of the targeted sequence along with an in-frame insertion of 24 bp sequence that resulted in the addition of 8 random amino acids (APTNTNYE) at the end of the protein. On western blots, *lon-1(ΔCTD)* mutant strains lack full-length LON-1 protein, and the V5-LON-1(ΔCTD) mutant protein is both stable and detectable (Figure 3B). We then examined the phenotypes of *lon-1(ΔCTD)* mutants with respect to BMP signaling and body size. As shown in Figure 3C and Table 1, both *jj373 jj536* and *jj373 jj537* mutants have a normal body size and exhibit no Susm phenotype. These results demonstrate that the CTD of LON-1 is dispensable for LON-1 function.

### LON-1 protein can bind cholesterol sulfate in vitro

Our finding that the CTD of LON-1 is dispensable and the fact that multiple previously identified missense mutations all affect residues in the CAP domain (MADUZIA *et al*. 2002; MORITA *et al*. 2002) suggest that the CAP domain of LON-1 must be critical for its function. Since multiple CAP domain proteins have been shown to bind sterol and/or fatty acids, including yeast Pry1 (CHOUDHARY AND SCHNEITER 2012; CHOUDHARY *et al*. 2014; DARWICHE *et al*. 2016; DARWICHE *et al*. 2017), mammalian CRISP2 (EL ATAB *et al*. 2022), plant PR-1 proteins *Solanum lycopersicum* (tomato) P14c and *Nicotiana tabacum* (tobacco) PR-1a (GAMIR *et al*. 2017), *Candida albicans* Rbe1p and Rbt4p (BANTEL *et al*. 2018), SmVAL4 and HpVAL-4 from parasitic nematodes *Schistosoma mansoni* and *Heligmosomoides polygyrus*, respectively (ASOJO 2011; KELLEHER *et al*. 2014; ASOJO *et al*. 2018), and since two *C. elegans* CAP domain proteins SCL-12 and SCL-13 also bind to cholesterol and function in the mobilization of cholesterol during dauer exit (SCHMEISSER *et al*. 2024), we tested whether LON-1 can also bind lipids in vitro.

We expressed and purified codon-optimized His-tagged LON-1 protein in *E. coli*, and used microscale thermophoresis to measure the affinity of LON-1 for cholesterol sulfate and palmitic acid. These binding assays showed that LON-1 binds cholesterol sulfate with a dissociation constant (*K_D_*) of ∼19 μM, while the *K_D_* of LON-1 binding to palmitic acid is ∼430 μM (Figures 4 and 5). These results indicate that LON-1 does not seem to bind fatty acid. Instead, it is capable of binding cholesterol sulfate in vitro. Although the affinity of LON-1 to cholesterol sulfate is lower than that observed for yeast Pry1 to free cholesterol (∼0.7 μM, (CHOUDHARY AND SCHNEITER 2012)), the binding is still in the micromolar range, as reported for other cholesterol-binding proteins (BUKIYA *et al*. 2011; SINGH *et al*. 2011; BUKIYA *et al*. 2017).

**Figure 4.**
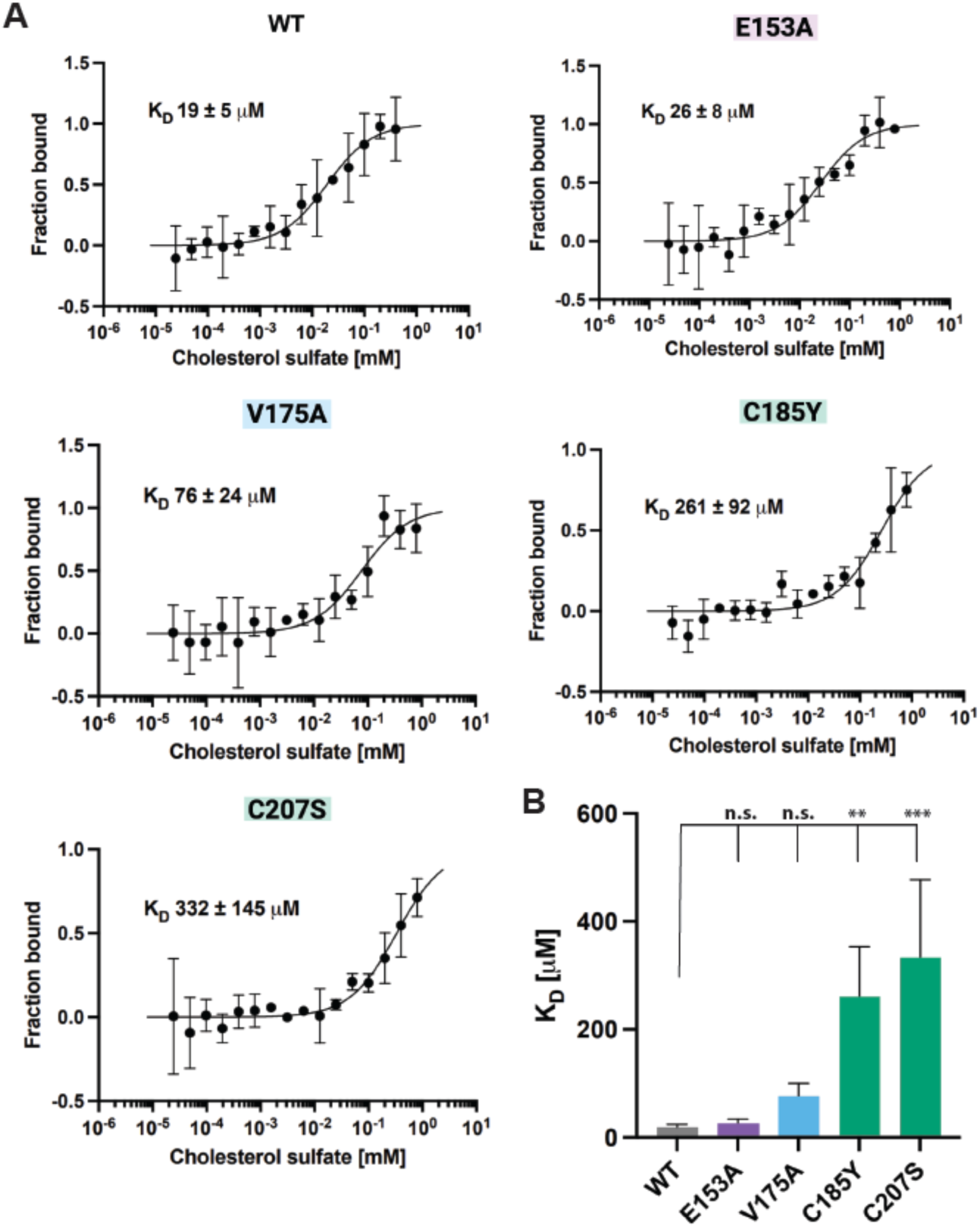
LON-1 binds sterols in vitro and mutations of conserved cysteine residues decrease the binding affinity. (A) The binding of cholesterol sulfate to wild-type and mutant LON-1 proteins measured using microscale thermophoresis (MST, see materials and methods). Cholesterol sulfate was titrated into 50 ng of fluorescently labelled proteins. K_D_ values are indicated and the values represent mean ± s. d. of three independent measurements. (B) Binding affinity (K_D_) as calculated in (A) were plotted for comparison. Asterisks indicate statistical significance (one-way ANOVA with Tukey’s post hoc test; \*\**p* <0.01; \*\*\**p* <0.001; n.s., not significant).

**Figure 5.**
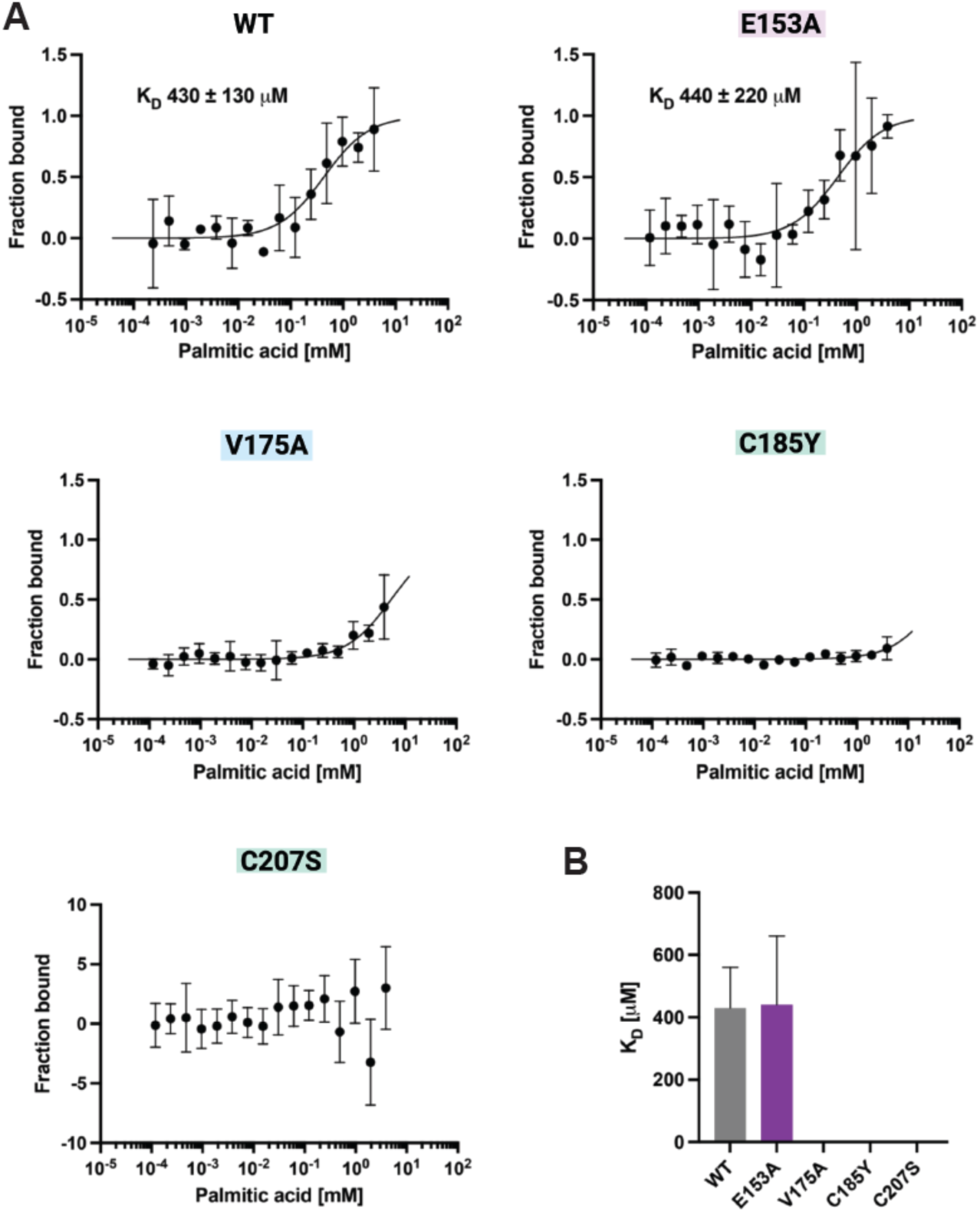
LON-1 binds palmitic acid in vitro at high micromolar concentrations. (A) Microscale thermophoresis (MST) dose response curves of fluorescently labelled proteins and fatty acids. Wild-type and mutant LON-1 proteins were tested for palmitic acid binding. The values represent mean ± s. d. of three independent measurements, and the dissociation constant (K_D_) are indicated. (B) K_D_ values obtained in (A) were plotted for comparison.

Previous studies on yeast Pry1 have identified several key residues important for Pry1 binding to cholesterol or fatty acid (CHOUDHARY AND SCHNEITER 2012; CHOUDHARY *et al*. 2014; DARWICHE *et al*. 2016; DARWICHE *et al*. 2017). Some of these residues are conserved in LON-1, including V175, whose equivalent in Pry1 is part of the putative fatty acid binding channel; E153 and H171, which represent two of the four residues that form a divalent cation binding pocket; as well as C185 and C207, which are predicted to form a disulfide bridge to stabilize the CAP domain and whose equivalent in Pry1 are required for sterol binding (CHOUDHARY AND SCHNEITER 2012). We therefore made point mutations in these specific residues and tested the mutant LON-1 proteins’ ability to bind cholesterol sulfate in vitro. As controls, we also conducted binding assays to palmitic acid.

As shown in Figure 4, the E153A mutation does not significantly affect LON-1’s ability to bind cholesterol sulfate. In contrast, LON-1 proteins carrying either the C185Y or the C207S mutation showed a greater than 10-fold reduction in their affinities to cholesterol sulfate (Figure 4A,B). LON-1 protein carrying the V175A mutation showed a somewhat reduced, but not statistically significant, affinity to cholesterol sulfate (Figure 4A,B). In addition, LON-1(V175A), LON-1(C185Y) and LON-1(C207S) mutant proteins also lost any weak binding to palmitic acid (Figure 5).

In summary, LON-1 protein can bind cholesterol, but not fatty acid, in vitro. Several highly conserved residues (C185 and C207) are important for LON-1’s binding to cholesterol.

### Key residues important for LON-1 cholesterol binding in vitro are important for LON-1 function in vivo

To begin to determine whether cholesterol binding of LON-1 is relevant for its function in vivo, we used CRISPR to generate several mutant lines, each carrying a mutation in one of the key residues as described above in the endogenous *lon-1* gene. To characterize these point mutant alleles, we measured the body size and the Susm penetrance of each mutant allele. The results are summarized in Figure 6. As was observed in the in vitro binding assay where LON-1(V175) did not have any obvious defect in binding to cholesterol sulfate, animals carrying the LON-1(V175A) mutation (*lon-1(kq175)*) did not exhibit any body size defect, though there was a slight, although statistically significant, increase in the Susm penetrance (5%, N=1244) compared to controls (0%, N=709) (Figure 6A,B). To exclude the possibility of *lon-1(kq175)* mutants showing a body length phenotype that was not detectable until later in development, we measured their body sizes at several additional time points when worms were well into adulthood and found no detectable changes in body size of *lon-1(kq175)* worms when compared to WT worms (Supplemental Figure 2). Since the V175A mutation targets the putative fatty acid binding channel, our in vitro and in vivo data suggest that LON-1 is likely not a fatty acid-binding protein.

**Figure 6.**
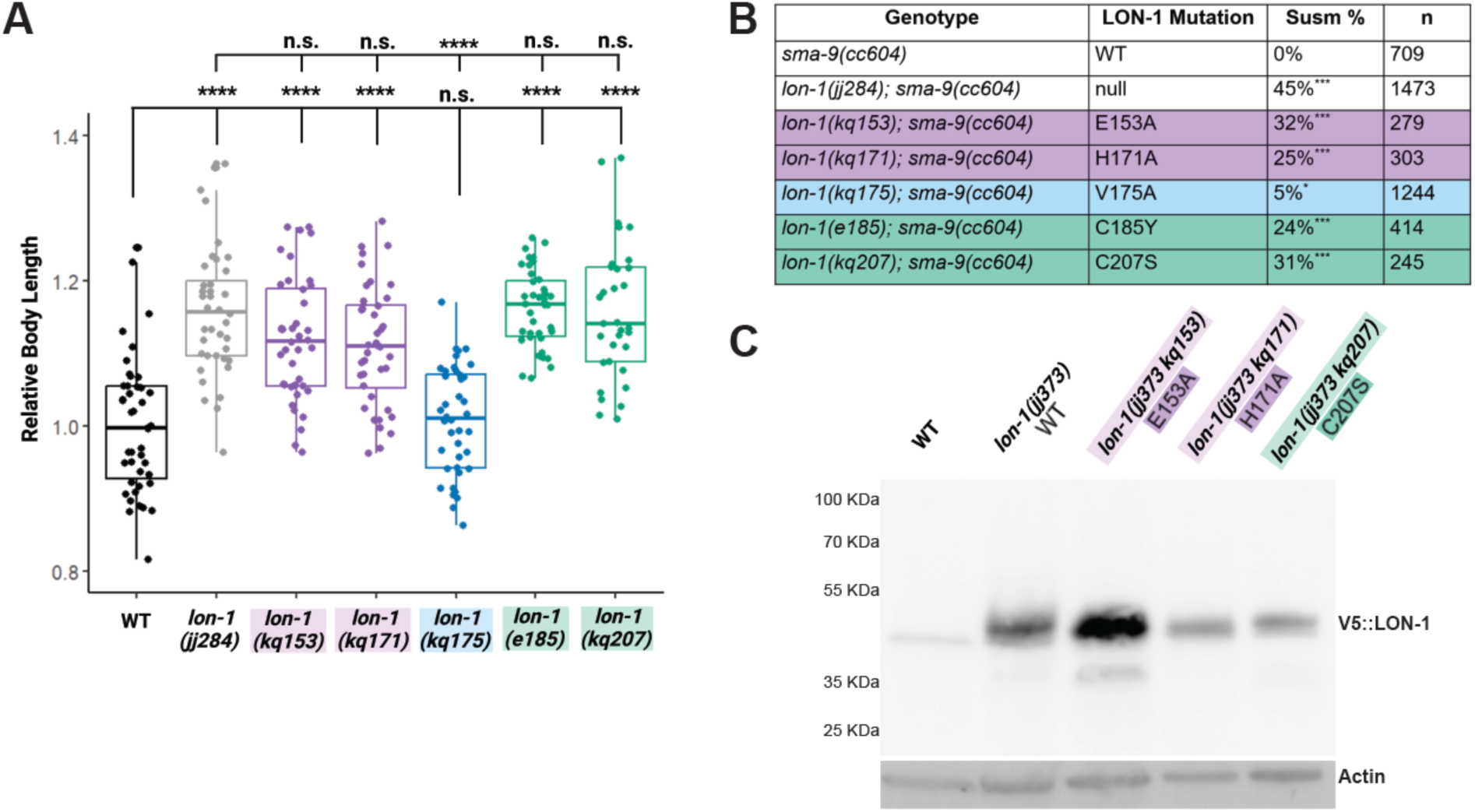
Point mutations in conserved residues in the CAP domain affect LON-1 function *in vivo*. (A) Relative body lengths of various strains at larval L4.1 stage, normalized to WT. Each dot represents one worm. **** *p*< 0.0001; n.s. not significant. Tested using an ANOVA with a Tukey HSD. (B) Table showing the Susm penetrance of various strains. Susm penetrance is measured as the percentage of 6 CC animals as scored by the *arIs37(secreted CC::GFP)* reporter. Statistical analysis was conducted by comparing double mutant lines with the *sma-9(cc604)* single mutants. *** *p* < 0.001; * *p* < 0.05 (unpaired two-tailed Student’s *t* test). (C) Western blot showing that mutant LON-1 proteins are detectable using anti-V5 antibodies (top). Each lane contains lysate from 100 gravid adult (GA) worms. Actin (bottom) was used as a loading control.

Unlike the V175A mutation, all other point mutations tested caused a strong body size phenotype. Mutant worms carrying the E153A (*lon-1(kq153)*), H171A (*lon-1(kq171)*), C185Y (*lon-1(e185)*) or C207S (*lon-1(kq207)*) point mutations are as long as *lon-1(jj284)* null worms (Figure 6A). Moreover, these four mutations also caused a Susm phenotype, with penetrance varying between 24% and 32%, slightly lower than the Susm penetrance of the *lon-1(jj284)* null allele (45%, Figure 6B).

To rule out the possibility that the point mutations that disrupted LON-1 function might have affected the expression or stability of mutant LON-1 proteins, we introduced several of the point mutations into the *lon-1(jj373[SP::V5::LON-1])* background, and conducted western blotting experiments using stage-matched adult worms. LON-1 proteins are detectable in all point mutants tested, which include E153A, H171A, and C207S, although LON-1 protein levels varied among the mutant strains (Figure 6C).

As described above, both LON-1(C185Y) and LON-1(C207S) mutant proteins exhibited significantly reduced affinity to cholesterol sulfate in vitro, and caused null-like phenotypes in body size and strong Susm phenotypes in vivo. C185 and C207 are predicted to mediate the formation of an intramolecular disulfide bond in the central beta sheet of the CAP domain (Figure 2F). It is likely that these two mutations affect the general structure of the protein, resulting in a loss-of-function phenotype. Nevertheless, LON-1(C207S) protein from mutant animals was detectable, albeit at lower levels than the WT protein, on western blots (Figure 6C). Intriguingly, mutating one of the equivalent residues in yeast Pry1 at C279 disrupted the sterol-binding ability of Pry1 (CHOUDHARY AND SCHNEITER 2012). A similar mutation in the tomato PR-1 protein P14c also affected the protein’s ability to bind sterol (GAMIR *et al*. 2017).

Both LON-1(E153A) and LON-1(H171A) mutant animals exhibit strong body size and Susm phenotypes (Figure 6A,B), yet LON-1(E153A) protein did not exhibit a decrease in binding affinity to cholesterol sulfate in vitro (Figure 4). Moreover, mutant LON-1(E153A) protein was consistently detected at an increased level compared to WT LON-1 protein (Figure 6C). E153 in LON-1 (E233 in yeast Pry1) is supposed to be one of four residues forming the ion binding tetrad that makes up the active site of CAP domain proteins (Figure 2D,E) (WANG *et al*. 2010; ASOJO 2011; MASON *et al*. 2014). Chelating Mg^2+^ prevented the sterol binding ability of Pry1 in vitro. However, an analogous E233A mutation in yeast Pry1 did not exhibit any defect in cholesterol binding in vitro, or prevent its ability to rescue the Pry1/2 double mutants in vivo (CHOUDHARY *et al*. 2014). Given that LON-1(E153A) did not show a reduction in sterol-binding affinity in vitro, it is possible that E153 is critical for sterol-independent functions of LON-1 in *C. elegans*.

## CONCLUSIONS

In summary, our in vitro and in vivo studies collectively suggest a role of LON-1 in regulating body size and BMP signaling via sterol binding. Since LON-1 is a secreted protein, it may carry sterols with it as it is transported through the secretory pathway to either deliver sterols to either the cellular membranes or to the extracellular matrix (ECM). Several members of the tetraspanin family, such as TSP-12, TSP-14 and TSP-21, are known regulators of the BMP signaling pathway and localize to both the cell surface and intracellular vesicles (LIU *et al*. 2015; WANG *et al*. 2017; LIU *et al*. 2020; LIU *et al*. 2022). Tetraspanins in mammals have been shown to form membrane microdomains known as tetraspanin enriched microdomains (TEMs) that contain cholesterol and glycosphingolipids (HEMLER 2005; YANEZ-MO *et al*. 2009). LON-1 could use an affinity for sterols to associate with the TEMs or transport sterols to the TEMs.

Alternatively, and not mutually exclusively, LON-1 may help transport/deliver sterols to the apical extracellular matrix (aECM). *C. elegans* worms are known to be coated with a cuticle that is composed of multiple layers of aECM components that include lipids and lipid-modified proteins (SUNDARAM AND PUJOL 2024). LON-1 is known to be produced by hypodermal cells (MADUZIA *et al*. 2002; MORITA *et al*. 2002), cells that are located directly underneath the aECM and are the major sites of production of aECM components (SUNDARAM AND PUJOL 2024). Such a role of LON-1 could help shape the worm body and regulate body size. Consistent with this idea, a number of cuticle collagen genes have been implicated in body size regulation, and reciprocal interactions between BMP signaling and cuticle collagens have been reported (NYSTROM *et al*. 2002; SUZUKI *et al*. 2002; YIN *et al*. 2015; MADAAN *et al*. 2018; LAKDAWALA *et al*. 2019; MADAAN *et al*. 2020; GOODMAN AND SAVAGE-DUNN 2022). Additionally, LON-1 may play a role in the mobilization of cholesterol during larval development, similar to two other *C. elegans* CAP proteins, SCL-12 and SCL-13, which can bind cholesterol and are involved in the mobilization of cholesterol upon dauer recovery (SCHMEISSER *et al*. 2024). In this case, free cholesterol released by SCL-12 and SCL-13-dependent internal deposits allows the activation of the mTORC1 pathway during the transition from quiescence to growth. Although mTOR signaling has not been implicated in the regulation of BMP signaling in *C. elegans*, BMP signaling is known to interact with insulin signaling to regulate lipid storage in *C. elegans* (CLARK *et al*. 2018; CLARK *et al*. 2021). A recent report identified genetic interactions between BMP signaling and AMPK and PI3K in regulating metabolism in *C. elegans* (VORA *et al*. 2022). LON-1 may play a role in mediating these interactions via its ability to bind sterol. Finally, LON-1 is likely to have sterol-independent functions as well, given the observations that LON-1(E153A) did not exhibit any sterol-binding defects in vitro yet caused severe body size and Susm phenotypes in vivo. E153 could be one of the key residues mediating interactions between LON-1 and its non-sterol binding partners. Ultimately, further investigation is needed to determine LON-1’s mechanism of action in both body size regulation and in BMP signal transduction.

## DATA AVAILABILITY STATEMENT

Strains and plasmids are available upon request. The authors affirm that all data necessary for confirming the conclusions of the article are present within the article, figures, and tables.

## ACKNOWLEDGEMENTS

We thank Josh Arribere, Dan Dickinson, Andy Fire, Bob Goldstein for plasmids, Erika Beyrent and Gunther Hollopeter for sharing CRISPR protocol, our late technician Herong Shi for generating the *lon-1* expression construct in S2 cells, and members of the Liu lab for helpful discussions and critical comments on the manuscript. S. M-A. was a Nexus Scholar of College of Arts and Sciences at Cornell University.

## SUPPLEMENTAL FIGURES

**Supplemental Figure 1. Alignment of *C. elegans* CAP domain proteins.**

(A) An unrooted tree generated using Clustal Omega (https://www.ebi.ac.uk/jdispatcher/msa/clustalo) to depict the relationship between LON-1 and other CAP domain proteins found in *C. elegans*. Protein sequences were acquired from UniprotKB with the search term “SCP containing protein” and limiting the taxonomy to *C. elegans*. The sequences of those that are shaded are used to generate the alignment shown in B. (B) Alignment was generated using Clustal Omega and formatted using Espript 3 (ROBERT AND GOUET 2014). The Hinge region is marked by a gray bar. The LON-1 CTD is highlighted in yellow. Solid red-shaded residues denote amino acids that are 100% identical across all proteins listed. Residues colored in red and framed in blue represent amino acids that are highly conserved. The numbers on top are based on the aa sequences of SCL-27. Parts of the N-termini of VAP-1, VAP-2, SCL-23 and F58E2.5 are not shown in this figure. VAP-2 has two CAP domains, which were named as VAP-2_N and VAP-2_C, respectively.

**Supplemental Figure 2. Relative body lengths of various *lon-1* point mutant strains at different developmental stages.**

Body length was normalized to that of young adult WT worms. Staged matched larval L4 stage worms were measured after approximately 48 hours of growth post bleach synchronization, young adult worms after approximately 60 hours of growth, and one day old adults after approximately 72 hours of growth. Each dot represents one worm. **** *p*< 0.0001; n.s. not significant. Tested using an ANOVA with a Tukey HSD.

